# Two decades of resurrection studies: What have we learned about contemporary evolution of plant species?

**DOI:** 10.1101/2025.07.27.667050

**Authors:** Lillie K. Pennington, Seema N. Sheth, Steven J. Franks, Jill T. Anderson, Elena Hamann

**Affiliations:** Genetics Department, University of Georgia, Athens, USA; Department of Plant and Microbial Biology, North Carolina State University, Raleigh, North Carolina, USA; Department of Biological Sciences, Fordham University, Bronx, New York, USA; Odum School of Ecology, University of Georgia, Athens, Georgia, USA; Department of Biology, Institute of Plant Ecology and Evolution, Heinrich-Heine University Düsseldorf, Düsseldorf, Germany

**Keywords:** Resurrection experiment, adaptability or adaptive potential, phenotypic plasticity

## Abstract

Global change has profoundly altered the eco-evolutionary trajectories of plant species. Longitudinal studies often document phenotypic shifts in response to climate change, such as earlier flowering in the spring, but it remains challenging to disentangle the contributions of phenotypic plasticity and adaptive evolution to shifted phenotypic distributions. The resurrection approach has emerged as a powerful method to study genetic and plastic responses to novel selection imposed by global change by contrasting ancestral and descendant lineages from the same population under common conditions. Here, we compiled a database of 52 resurrection studies to examine key hypotheses about plant evolutionary responses to global change using a meta-analysis (40 of the studies) and quantitative review (all 52 studies). We found evidence for rapid, contemporary evolution, which often appeared adaptive, in over half of the cases, including some of the fastest cases of evolution in natural populations ever observed. Annual plants evolved earlier reproduction, and leaf economic traits associated with stress escape strategies. We also found evolution of increased plasticity for annual plants in phenology and physiology traits, and a reduction of plasticity in traits related to the leaf economic spectrum. We found less evidence for evolution in perennial species. Overall, our findings demonstrate the key role of drought escape in plant responses to a warming world. However, the lack of evolution in other traits and species indicates that constraints may dampen evolutionary responses in some scenarios. Our review also suggests promising avenues of future research for resurrection studies.

## Introduction

Within the last 40 years, a burgeoning literature has substantially deepened our understanding of contemporary evolution in natural populations (Endler, 1986; Grant and Grant, 2014; Thompson, 2013; Kingsolver and Buckley, 2017). This work has revealed that rapid evolutionary changes can occur within just a few generations (Grant and Grant, 2002; Franks et al., 2007; Schoener, 2011), and evolutionary processes have the potential to influence population persistence on relatively short timescales (Carroll et al., 2007; DeMarche et al., 2019; Anderson et al., 2025). Despite these enormous advances, it remains challenging to demonstrate that adaptive evolution has occurred in a given natural population. This task has become more pressing in an era of rapid environmental change, as adaptive responses to novel selection could enable species to persist in their current ranges (Ellner et al., 2011; Chevin and Bridle, 2025). One recent methodological advancement that addresses these issues is the direct comparison of traits expressed by ancestor and descendant lineages from the same location under common conditions (a ‘resurrection study,’ Franks et al., 2007, 2018). The resurrection study is a powerful approach for quantifying the extent and directionality of contemporary evolution (Franks et al., 2007, 2018). Here, we performed a quantitative literature review and a meta-analysis of resurrection studies to characterize insights into evolution in wild populations gained by this approach.

The meta-analysis framework can illuminate major evolutionary patterns that have emerged across resurrection studies despite the substantial variation in evolutionary responses among species that differ in life histories and mating systems, populations that differ in genetic diversity and connectivity, traits that differ in heritability and plasticity, and regions that differ in the drivers and rate of environmental change. For example, species with fast generation times and high standing genetic variation may adapt more rapidly to strong selection imposed by contemporary climate change (Hamann et al., 2018), whereas long generation times, low genetic diversity and genetic trade-offs across traits could constrain adaptation in others (Etterson and Shaw, 2001; Jarrold et al., 2019; Moran, 2020). As such, it follows that annual species may have faster evolutionary rates, in absolute time, than perennial species because of the shorter generation times of annuals. Ultimately, we seek to understand how variation in life-history traits, ecological strategies, population size, mating systems, standing genetic variation, and intra-specific genetic differentiation (Thompson et al., 2023) could contribute to differences in the adaptive potential of populations confronting novel environmental conditions (Etterson and Shaw, 2001).

A synthesis of the resurrection study literature can also examine the diversity of evolutionary responses to stress, especially given that species with different life history strategies may have contrasting evolutionary responses to the same abiotic stress. The direction and type of evolutionary responses could be driven by resource acquisition strategies, mirroring several classical theoretical frameworks (Fig. 1). Coordinated suites of traits often evolve in response to competition, resource limitation, and periodic biomass destruction via environmental disturbance (Grime, 1977, 1974; Reich, 2014; Pierce et al., 2017). These trait syndromes have been captured elegantly along the Leaf Economics Spectrum (LES) (Wright et al., 2004). Specifically, stress resistant plants tend to display slow growth, long life-spans, and a resource-conservative strategy, characterized by high leaf dry matter content, low leaf area, low specific leaf area and reduced leaf nitrogen content. In contrast, other species often express resource-acquisitive traits at the fast end of the leaf economic spectrum (Fig. 1). For example, frequent and severe droughts, which are increasing with climate change (IPCC, 2023; Kornhuber et al., 2024), may favor the evolution of drought escape strategies in annual species and drought avoidance or tolerance strategies in perennials (Kooyers, 2015). Specifically, in response to drought stress, annual species often evolve earlier flowering (Franks et al., 2007; Vigouroux et al., 2011; Nevo et al., 2012; Thomann et al., 2015; Hamann et al., 2018; Rauschkolb et al., 2022a), earlier seed emergence (Dickman et al., 2019), and accelerated growth (Franks, 2011; Rauschkolb et al., 2022a). These phenotypic changes are consistent with a drought escape strategy characterized by rapid growth and development (Wright et al., 2004). In contrast, we predict perennials that more often display a stress resistant and resource-conservative strategy will evolve toward suites of traits enabling drought tolerance/avoidance. The resource-conservative strategy will be accompanied with delayed flowering due to slow growth, assimilation rate and stomatal conductance, and high water-use efficiency (Fig. 1). Indeed, a resurrection study with a perennial herb documented the evolution of drought avoidance, where plants delayed flowering and increased water-use efficiency (Anstett et al., 2021)(Fig. 1). However, the consistency of these responses remains unclear.

**Fig. 1:**
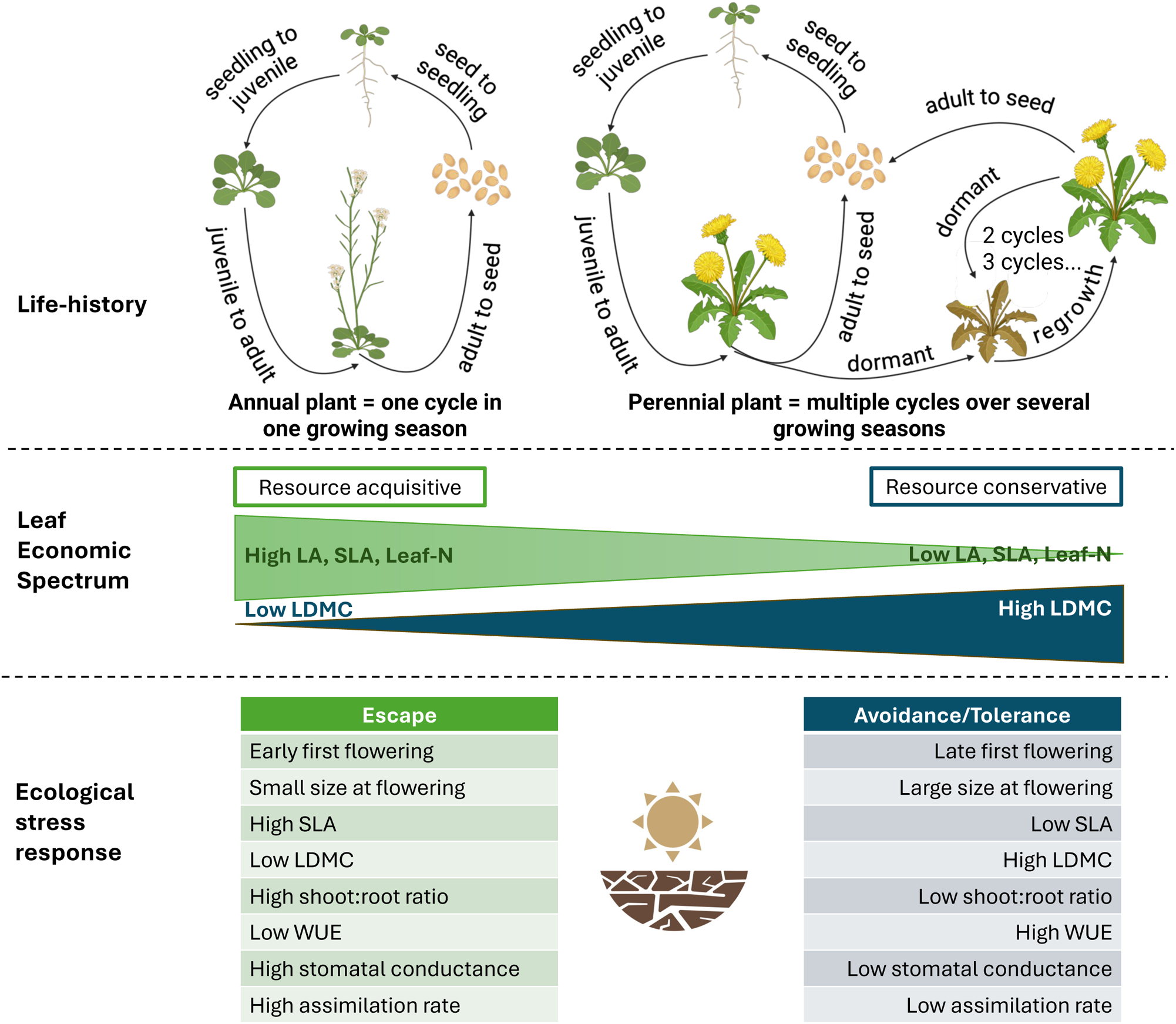
Integrating across several classical theoretical frameworks to predict plant responses to drought stress based on their life-history traits, resource-use strategy as captured on the Leaf Economics Spectrum, and their ecological classification. We predict that annual plants, which often display a resource-acquisitive strategy, are more likely to respond to drought stress via an escape strategy characterized by fast growth and early flowering, small size at flowering, high SLA, low LDMC, and accompanied by low WUE, high stomatal conductance and assimilation rates. In contrast, perennial plants that often display a stress resistant and resource-conservative strategy to ensure their persistence may instead evolve towards drought tolerance or avoidance strategies and display slow growth, delayed flowering, large size at flowering, low SLA, high LDMC, and high WUE with low stomatal conductance and assimilation rates. Figure generated with BioRender.

In addition to quantifying the extent of trait evolution, resurrection studies that expose descendant and ancestral lineages to multiple environmental conditions can evaluate the evolution of trait plasticity (Sultan et al., 2013; O’Hara et al., 2016; Hamann et al., 2018; Lambrecht et al., 2020; Johnson et al., 2022a, 2022b; Rauschkolb et al., 2022b, 2022a; Anstett et al., 2021; Christie et al., 2023; Spear et al., 2023; White et al., 2023; Branch et al., 2024; Spurlin and Lambrecht, 2024; and see Matesanz et al., 2010). When individuals experience multiple environments across their lifespan or when their progeny disperse into habitats that differ from those of the parents, trait plasticity could confer increased fitness and represent an adaptive response to spatial and temporal variation in environmental conditions (Moran, 1992; Alpert and Simms, 2002; Baythavong, 2011; Anderson et al., 2012). Indeed, plasticity may evolve readily in response to spatial and temporal variation in conditions (Murren et al., 2014). Increasing environmental variability under climate change (Lázaro-Nogal et al., IPCC 2023, 2015; Kornhuber et al., 2024) could impose selection on norms of reaction for various traits (e.g. Jentsch et al., 2007; Kingsolver and Buckley, 2017; Matesanz et al., 2010; Stotz et al., 2025), and adaptive plasticity could mitigate the immediate negative fitness effects of climate change (Matesanz et al., 2010; Nicotra et al., 2010; Shaw and Etterson, 2012; Franks et al., 2014). However, not all populations maintain genetic variation for trait plasticity, not all traits are plasticity, genetic correlations could constrain the evolution of increased plasticity (Murren et al., 2014; Oostra et al., 2018), and climate change is not uniform across the globe (IPCC 2023). Furthermore, extreme events, such as heat waves are projected to increase under climate change (van der Wiel and Bintanja, 2021; Cai et al., 2022; Kornhuber et al., 2024), and we know little about how rapid year-to-year shifts in selection could influence the evolution of plasticity (Chevin and Hoffmann, 2017). Thus, it remains unclear whether changing climates will favor the evolution of plasticity or whether plasticity will be sufficient to maintain fitness especially under novel climatic extremes (Duputié et al., 2015; Chevin and Hoffmann, 2017; Kingsolver and Buckley, 2017; Arnold et al., 2019; Stotz et al., 2025). Resurrection studies conducted in multiple ecologically-relevant environments are uniquely situated to investigate genetic shifts in plasticity in response to factors associated with global change, and when paired with selection analyses, these studies can evaluate the adaptive nature of plasticity.

In 2018, Franks and colleagues reviewed resurrection studies that revived stored ancestral propagules along with contemporary descendants to examine evolutionary changes in natural plant populations under common conditions. At that time, 12 studies had implemented a resurrection approach, all of which reported rapid, contemporary evolution in one or more traits. Furthermore, fitness increased across generations for most of these instances, revealing rapid adaptation. Franks et al. (2018) noted that inferences from resurrection studies could potentially be biased due to maternal effects (Roach and Wulff, 1987), in which adult traits of offspring are influenced by the environmental conditions of the maternal plant, to storage conditions effects, in which adult offspring traits are influenced by the conditions that the stored ancestral seeds experienced (Franks et al., 2018), or by the invisible fraction effect, in which there is non-random mortality during storage (Weis, 2018; Franks et al., 2019). These issues are minimized when there is a ‘refresher generation’, where initial collections of both ancestors and descendants are grown in common conditions, as this reduces maternal and storage conditions effects, and when there are very high germination rates for both ancestral and descendant seeds (Franks et al., 2018). While the majority of studies in our database included a refresher generation, many did not, and this could potentially influence results. Studies also differed in their use of experimental manipulations: in the initial review, 5 studies out of 12 grew ancestors and descendants under common, benign conditions, which can detect genetic change in traits, but cannot address whether these evolutionary changes were adaptive nor can it quantify evolution of reaction norms. Finally, few studies estimated quantitative genetic variation or heritability, limiting our understanding of the adaptive potential of ancestral vs. descendant populations.

Here, we conduct the first formal meta-analysis of resurrection studies to evaluate four key hypotheses about plant evolutionary responses to novel environments. With this meta-analysis, we first aim to test whether annual plants respond more rapidly to environmental change than perennials (H1), whether annuals and perennials have different patterns of evolution across broad categories of traits generally (H2a), and whether, in the context of increasing drought stress, annual plants will evolve traits associated with drought escape, whereas perennial plants will develop traits linked to drought avoidance or tolerance (H2b). In response to increased environmental variability associated with climate change (Moran, 1992; Baythavong, 2011; Anderson et al., 2012), plant populations will evolve increased plasticity (H3). We had stringent selection criteria for our formal meta-analysis, but we also conducted a quantitative literature review to provide a broader view of resurrection studies and to highlight general trends in the published literature, including studies not retained in our meta-analysis.

## Methods

### Meta-analysis search criteria

We conducted a phylogenetically-corrected meta-analysis to examine evolutionary responses to contemporary climate change. In October 2023, January 2024, and April 2025, we searched for resurrection studies on Web of Science using the terms “resurrection approach” AND “plant” and “resurrection study” AND “plant” with the aim of extracting phenotypic data and fitness components from all plant resurrection experiments published to date. We supplemented this search with a Google Scholar query using the terms ‘resurrection study’ AND ‘plant’ and used the ‘search within citing articles’ feature to identify publications citing foundational resurrection studies (Franks et al., 2007, 2008) and the initial literature review (Franks et al., 2018,). We screened the titles and abstracts of 52 studies identified through these searches to identify suitable publications based on these inclusion criteria: (1) the study contrasted ancestral and descendant lineages in an experimental framework, either in the field or controlled conditions (e.g., greenhouse or growth chamber) and measured traits and/or fitness for at least one ancestral and descendant generation from at least one natural population; (2) efforts were made to sample seeds or propagules from the same natural populations in both ancestral and descendant generations; (3) data were available for both generations, either within the publication (in figures or tables), through online repositories, communications with authors, or in supplemental material such that we could download sample size, means, and either standard deviations or standard errors or extract these parameters from figures with WebPlotDigitizer version 4.8 (Rohatgi, 2024); and (4) include direct measurements of phenotypes and/or fitness. This rigorous screening process ensured that only studies with sufficient and standardized data on evolutionary responses to climate change in plants were included in our meta-analysis. Out of the 52 resurrection studies we identified, 40 studies (76%) met these criteria. As only five studies of three species presented genomic sequence differences across generations, we did not consider genomic data in the formal meta-analysis (but see Discussion). We also omitted a study (Wooliver et al., 2020) that focused on the evolution of thermal performance curve parameters because these are function-valued traits estimated from statistical models rather than directly measured traits.

### Qualitative data extraction

In addition to our formal meta-analysis, we conducted a comprehensive overview of all resurrection studies by compiling information from all resurrection studies that reported phenotypic traits, fitness components, and genomic sequence data (Table S1). For each study, we recorded the research questions, aims, and hypotheses, along with key elements of the work such as: the geographic extent of seed/propagule sampling for ancestral and descendant genotypes; the number of maternal lines and individuals sown; the experimental design and treatments applied; and the traits and fitness components measured (Table S1). We summarized key results, noting which traits were reported to have shown evidence of evolution, whether evolutionary changes were discussed or tested as adaptive, what the adaptive response was, if evolution in plasticity was observed or discussed, whether heritability was assessed, the reported rate of evolution (if calculated), and the putative driver of evolutionary change (Table S1).

### Quantitative data extraction

From each publication that met our inclusion criteria, we extracted trait and fitness data for both the ancestor and descendent generations by calculating the mean, standard deviation, and sample size using the summarySE function in the R package Rmisc (Hope RM 2022) for each population as necessary from raw archived data. We obtained complete datasets for almost all studies from data repositories and communication with authors. For two studies, we used WebPlotDigitizer version 4.8 (Rohatgi, 2024) to extract data from figures, and we extracted data provided in a supplementary table for one study. For studies which contrasted ancestral seeds with descendant seeds collected across multiple years (N=3), we included only the earliest and latest years in which more than one individual was measured in the meta-analysis.

To examine trends across studies, we categorized traits into eight groups: resource allocation, fitness components, floral traits, leaf economic spectrum traits, other leaf traits, phenology, physiology, and plant size (Table 1; see also Siefert et al., 2015; Griffin-Nolan et al., 2018; Lemmen et al., 2019; Pennington et al., 2021; Boyd et al., 2022). Very few studies reported binary fitness components (e.g., survival, percent flowering, percent emergence, Table S1) and were therefore not included, as binary data requires the use of odds ratios as an effect size, which cannot easily be combined with the Hedge’s g, which is an effect size for continuous data. We recorded the number of intervening years between the ancestor and descendant generations, whether or not a refresher generation was grown before the experiment, whether the species was native or invasive to the region where seeds were collected, the mating system (primarily selfing or primarily outcrossing), the life history strategy (annual or perennial), the experimental setting (field, greenhouse, growth chamber), and which manipulations (if any) were applied. Furthermore, we aimed to extract data on generation times. However, we found uneven representation of trait categories across these qualitative variables; the current studies are biased towards native, annual plants in lab settings (Table S1, Fig S2 & S3). Furthermore, data on mating systems and generation times were often missing. In cases where data were available, many species operated along a mating system spectrum—rather than being strictly selfing or obligately outcrossing. Therefore, we focused our analyses on trait category and life history strategy, which had the most comprehensive representation across the dataset.

**Table 1.** Trait categories used in the current meta-analysis with definitions and the reported traits that were sorted into each category.

| Category | Definition | Traits reported in publications |
| --- | --- | --- |
| Leaf economic spectrum (LES) | Global framework that captures trade-offs in leaf investment and resource use efficiency | Specific leaf area (SLA), leaf dry matter content (LDMC), leaf area (LA) |
| Leaf traits | Structural and anatomical characteristics that influence gas exchange, water retention, and defense, but fall outside the conventional leaf economic spectrum framework. | Leaf succulence, leaf length, leaf width, stomatal density, trichome density, secondary compound concentration, leaf mass, guard cell length |
| Physiology | Measurements of metabolic function, reflecting resource processing and energy acquisition | Carbon assimilation, stomatal conductance, carboxylation rate, respiration, photosynthetic rate, osmotic potential, fluorescence, water use efficiency (WUE) |
| Phenology | Developmental timing | Time to first flower, time to emergence, mean flowering time, flower duration |
| Fitness | Proxies for the genetic contribution to the next generation | Number of flowers, pollen count, seed number, pollinator/selfed ratio, seed mass, seed count, achene number, ramet number, reproductive biomass, achene size, achene mass, seed/ovule ratio |
| Plant size and growth | Measurements of plant size and growth rates | Growth rate, biomass, leaf number, root biomass, height, plant width, diameter, height at flowering, node of first flower, aboveground biomass, belowground biomass, branch number, stem density, rosette size |
| Floral traits | Characteristics of individual flowers or inflorescences | Anther-stigma distance, flower size, stigma length, nectar brix nutrition, flower display, flower duration, flower color, flower diameter |
| Resource allocation | Traits associated with allocation of resources | Root/shoot ratio, reproductive allocation, belowground distribution |

We calculated rates of evolution, in haldanes, following the formula given in Hendry and Kinnison (1999), which gives evolutionary rates in standard deviation units:

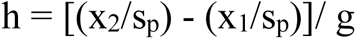

where h= haldanes (the rate of evolution),

x_2_= the mean trait value of descendants,

x_1_= the mean trait value of ancestors,

s_p_ = the pooled standard deviation

g = the number of generations.

As this formula includes the number of generations, and we did not know precisely the number of generations in the studies of perennials, we limited this analysis to the annual plants in our dataset (using the number of intervening years between ancestors and descendants as an approximation of the number of generations; see Table S1).

### Phylogenetic correlation matrix

To correct the meta-analysis for phylogeny, we constructed a phylogenetic correlation matrix using data from the Open Source Tree of Life (OpenTree et al) via the rotl package ver. 3.1.0 (Michonneau et al 2016) in R and the ape package ver. 5.7.1 (Paradis and Schliep 2019). When necessary due to taxonomic name changes (e.g., *Mimulus* sp. are referred to as *Erythranthe* in the TOL), we updated the reported species name to match the name in the Tree of Life; six species had their names updated. We then produced our tree using the tol_induced_subtree function in rotl and calculated branch lengths using the compute.brlen function in the ape package (Paradis and Schliep 2019). Finally, we computed a phylogenetic correlation matrix for use in our meta-analysis using the vcv function in the ape package (Paradis and Schliep 2019), which we included as a random effect in all meta-analysis described below.

### Meta-analysis

We performed all meta-analyses in the R package metafor ver. 4.8-0 (Viechtbauer, 2010). We computed the standardized mean difference effect size (Hedges’ *g*) (Hedges, 1981) and variance with the escalc function in metafor. Our calculation of effect size compares ancestors and descendants, with positive numbers indicating that the descendant generation had larger trait or fitness values than the ancestral generation.

Our dataset combines several leaf economic traits that reflect drought escape in slightly different ways. For example, high specific leaf area (SLA) and low leaf dry matter content (LDMC) signify drought escape whereas low SLA and high LDMC indicate a drought avoidance or tolerance strategy (Kooyers, 2015). To ensure that all traits within a category were biologically meaningful, we converted specific leaf area to its inverse (leaf mass area) by multiplying the effect size for specific leaf area by negative one (-1). This change meant that the directionality of this trait matched expectations for drought escape (i.e., low leaf mass area and LDMC signal drought escape). Further, three studies measured “relative water content” or “foliar water content” as leaf wet mass/dry mass — for these instances, we categorized them as LDMC but again multiplied by -1 for the sign to match directionality of LDMC, which is measured as leaf dry mass/wet mass.

Studies varied substantially in the number of years that elapsed between the ancestral and descendant generations. To account for this variation, we divided each effect size by the number of intervening years between the generations, resulting in effect sizes scaled by time elapsed between the ancestor and descendent generations. We adjusted variance in the scaled rate effect size by dividing the variance in the overall effect size by the square of the number of intervening years. We highlight that very few studies of perennials included information about the exact generation time of the focal species, nor was it possible to extract these data for all species from other sources; therefore, we were not able to correct our effect sizes for the number of generations separating ancestors and descendants. However, we did include annual vs. life history strategy as a moderator in the analyses to examine explicitly whether annuals and perennials have different evolutionary responses, in absolute rather than in generation time, to novel conditions. We used the rma.mv function in metafor to analyze the scaled effect size as a function of a series of moderators (fixed effects), including life history (perennial vs. annual), trait category, and their interactions. All analyses included study identifier, species name, population identifier, and the phylogenetic correction as random effects. For studies with only a single population, we assigned a study-specific stand-in population identity. We conducted three separate statistical tests using our dataset to address our hypotheses. To account for testing multiple hypotheses with one dataset, we set our alpha to 0.017 (=0.05/3 tests).

#### Analysis 1

To examine whether annual plants respond more rapidly to environmental change than perennials (H1) and if annual and perennial plants have different evolutionary outcomes for different traits (H2a), we analyzed the rate of evolution (effect size scaled by number of intervening years) as a function of life history, trait category, and their interaction.

#### Analysis 2

To test whether annual species evolved a drought escape strategy and perennials evolved drought tolerance (H2b), we examined the subset of studies that imposed a drought treatment. We then analyzed the rate of evolution as a function of life history, trait category, and their interaction.

#### Analysis 3

To test whether descendants have evolved higher plasticity (H3), we again examined the subset of studies that imposed a drought treatment, in this case because this was the only treatment that was imposed in enough studies (n = 19) for sufficient replication. We calculated the relative distance plasticity index (RDPI) for each generation and trait as the absolute difference between the means of the control and experimental treatment, divided by the sum of those means (Valladares et al., 2006). We then calculated Hedges’ g using the RDPI values for each generation and trait as the descendant RDPI minus the ancestor RDPI, resulting in an effect size which relates to the evolution of plasticity. We again scaled this effect size by the number of intervening years, resulting in a rate of change in plasticity. We analyzed this rate as a function of life history, trait category, and their interaction.

## Results

### Quantitative literature review

Our full literature search returned 52 resurrection studies, of which 96.2% detected phenotypic evolution, with descendants differing significantly from ancestors in 123 of the 243 measured traits (51%). Only two studies (3.8%) reported no significant change between the ancestor and descendant generations in any of the traits measured (Table S1). We found 17 studies that measured plasticity, 7 that estimated heritability, and 4 that reported the rate of evolution in haldanes. The number of populations included in each study ranged from 1-60, with an average of 10 populations per study. Adaptive responses to climate change likely vary across the range of a species (Eckert et al., 2008; Pironon et al., 2017; Pennington et al., 2021); however, only five resurrection experiments used populations collected from across the full geographic distribution of the species (Kooyers, 2015; Dickman et al., 2019; Vtipil and Sheth, 2020; Wooliver et al., 2020; Anstett et al., 2021; Branch et al., 2024) and an additional two studies collected from many populations but only within a single country (Nevo et al., 2012; Al-Hajaj et al., 2022). We found mixed reporting on mating system, with 16 studies not reporting the mating system of the focal species. Five studies did not indicate whether the species was native to the location where source populations were identified– when not identified, we searched the literature to add this information to our table. The number of maternal families included ranged from a minimum of 8-385 per population, with the total number of individuals ranging from 20-6912. Thirteen studies detected evolution in plasticity, with nine reporting gains in plasticity in at least one trait and six reporting losses (Table S1).

In calculating rates of evolution in the annual plants in the dataset, we found evidence for rapid evolution, but also substantial variation within and between trait categories. Across all traits and species, the average absolute value of the evolutionary rate was 0.090 haldanes (n = 1181, SE = 0.0038). However, there is also substantial variation around this average rate, with cases ranging up to 1.65 haldanes. When examined by trait category, leaf economic spectrum traits and phenology had the fastest rates, and allocation, floral traits and physiology the slowest (Table 2, Fig. S4). In terms of directionality, we found several trends, including, for example, that overall fitness evolved to be greater while phenology evolved to be earlier (Table 2).

**Table 2.**
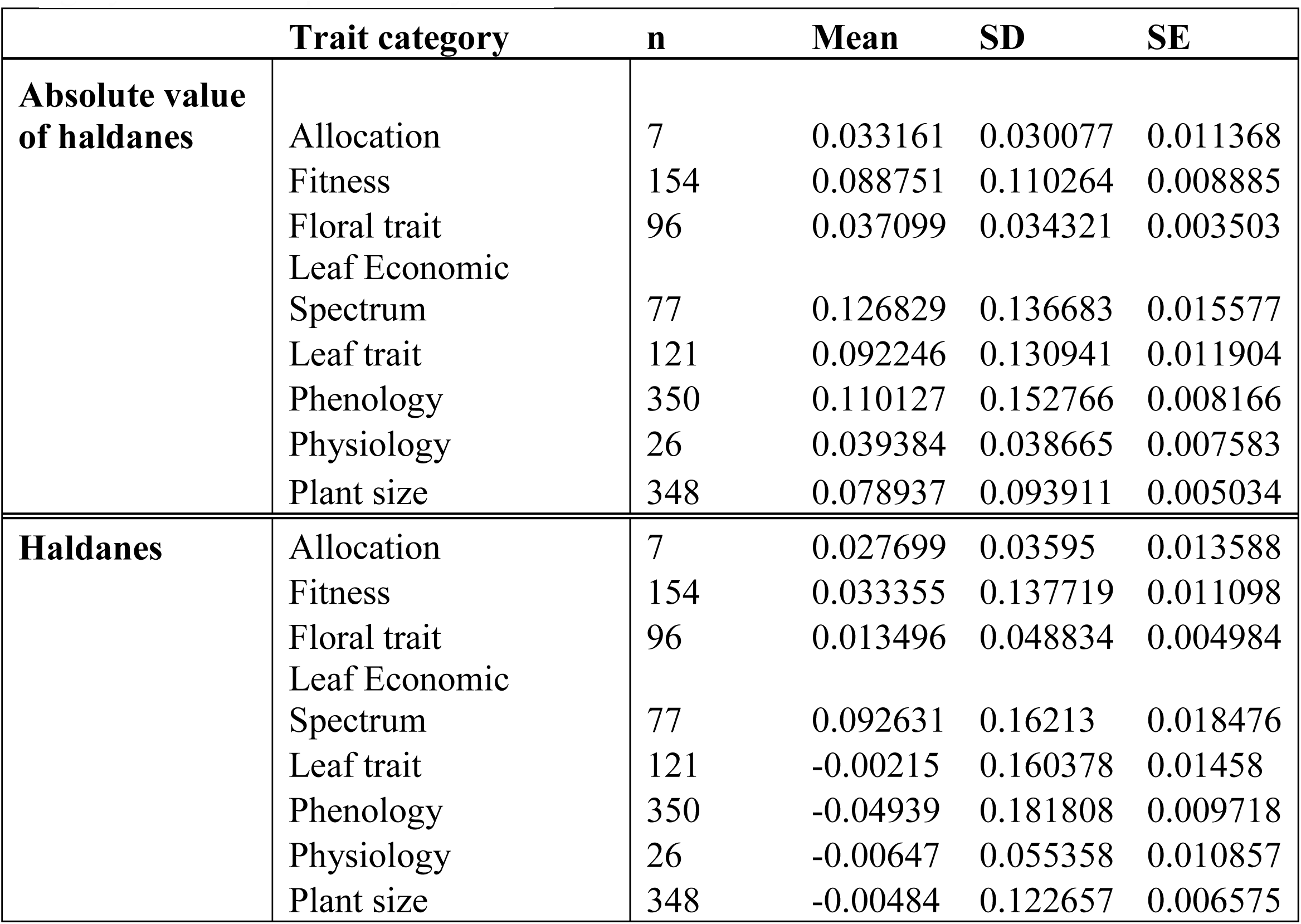
Rate of evolution reported in absolute value of haldanes and haldanes, for each trait category for annual species only.

We only found seven published resurrection studies with genomic or genetic data (Table S1), Of these, three experiments paired genomic and phenotypic data to examine genetic changes between the ancestral and descendant generations. Studies have found changes in gene frequency in genes related to drought stress and flowering time (Rhoné et al., 2010; Nevo et al., 2012; Franks et al., 2016), as well as upregulation of those genes (Hamann et al., 2021c).

### Formal Meta-analysis Overview

Of the 52 studies that we identified for the literature review, five were excluded from the meta-analysis as they reported exclusively genomic data, four were excluded for studying ancestral and descendant populations that were not geographically the same, two for data unavailability, one was excluded for having unclear sample sizes, and one for reporting change in statistical parameters rather than raw phenotypes. Thus, 40 studies met the inclusion criteria for the meta-analyses. In sum, these 40 studies included 21 orders, 28 families, and 87 species of flowering plants (phylogeny Fig. S1) in 5 continents, and resulted in 1761 effect size observations. The dataset had 582 observations from perennial species and 1179 from annual species (Fig. 3c). The mean number of intervening years between the ancestor and descendant populations was 13.2 years (standard deviation: SD= 13) with a range of 2-53 years. In the full dataset, 635 observations were from experimental manipulations of factors such as drought and herbivory while 1126 were from benign, well-watered conditions that were designed to avoid a stress response. A refresher generation was used in 70% of included studies (Table S1).

**Figure 2.**
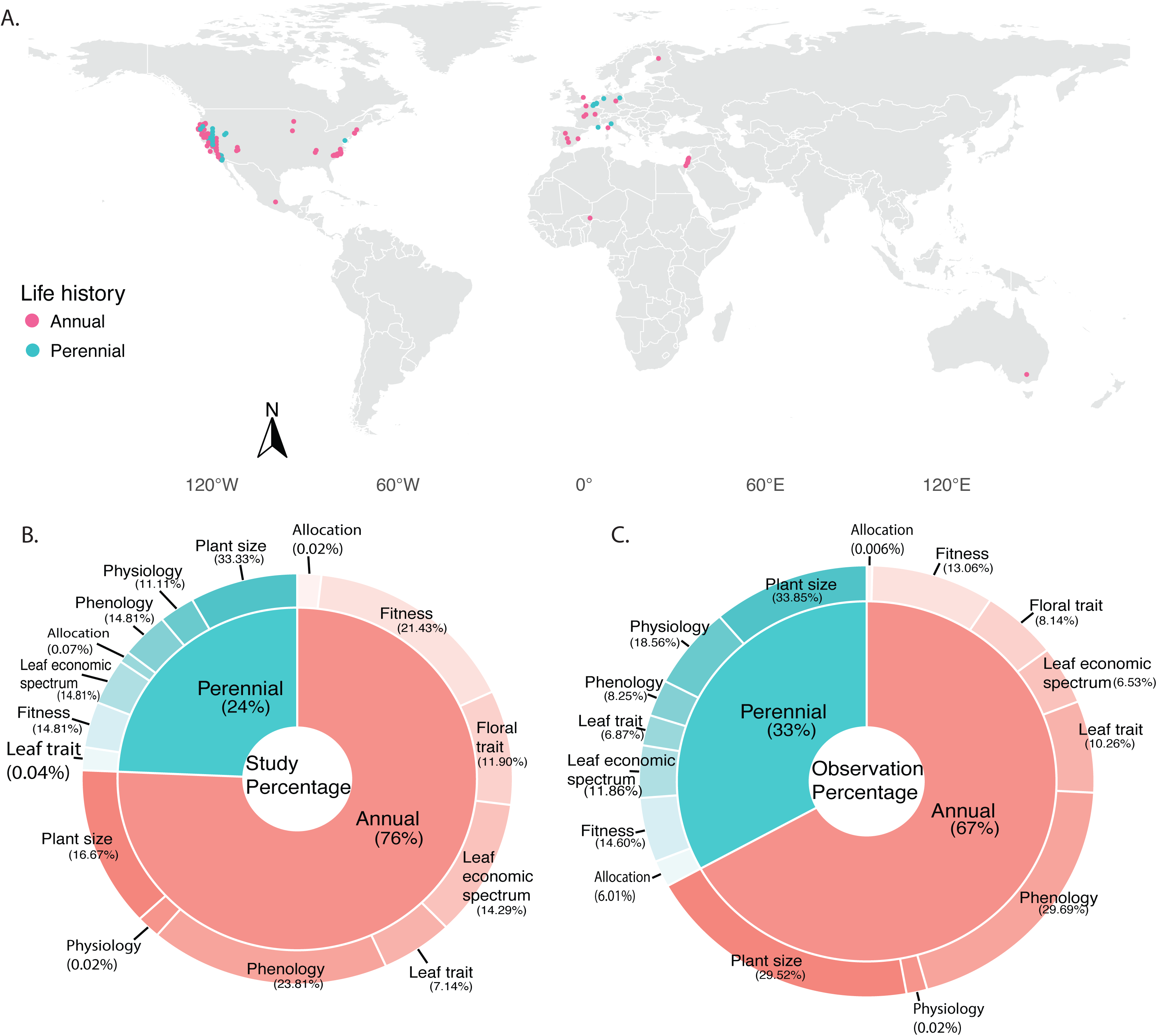
(A) World map showing distribution of study sites. (B) Proportion of studies partitioned by life history strategies and associated trait categories in the dataset. (C) Proportion of observations partitioned by life history strategies and associated trait categories in the dataset. For B and C, the inner circle represents the proportion of studies classified by life history strategies. The outer circle shows the distribution of trait categories within each life history strategy, with percentages indicating the relative contribution of each trait.

**Figure 3.**
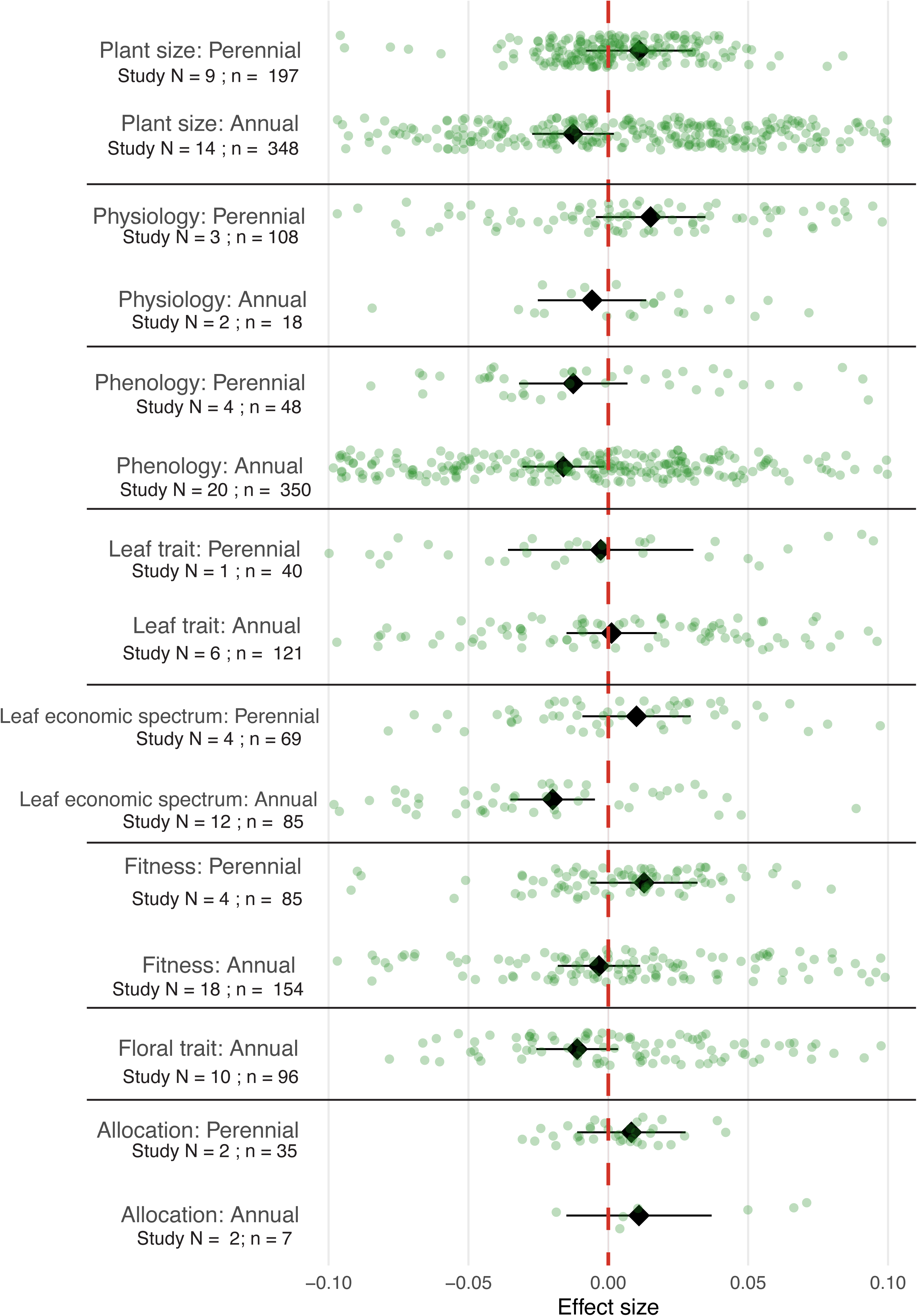
Interaction effects of trait category and life history on evolutionary rates. Effect sizes are represented by black diamonds, with horizontal lines indicating 95% confidence intervals. Positive values indicate that descendants have larger trait values than ancestors. Green points represent raw effect sizes. The x-axis is truncated to highlight patterns more clearly, full data is available as supplementary material. The overlap of confidence intervals with zero for some categories suggests non-significant effects in those cases. Study N indicates the number of studies that included each trait category; n is the number of observations included in the analysis. Dashed red vertical lines represent an effect size of 0.

### Hypotheses 1 and 2a: Annual plants respond more rapidly to environmental change than perennials, and they differ from perennials in the directionality of trait evolution

Our meta-analysis found significant effects of trait category (n = 1746, QM = 133.12, df = 8, p-val < .0001) and the interaction between trait category and life history (n = 1746, QM=133.12, df=6, p<0.0001, Table S2), indicating that the magnitude and directionality of evolutionary shifts differs for annual and perennial plant species across traits. Specifically, annual descendants had lower values for traits related to the leaf economic spectrum (LES), reflecting the evolution of a drought escape strategy (low leaf dry matter content, high specific leaf area, low leaf mass area) as well as earlier phenology than annual ancestors, with floral traits and plant size trending lower. Floral traits were not quantified in perennial species in our dataset. Perennial descendants did not significantly differ from ancestors, though phenology also trended earlier and other trait values like physiology, LES, fitness, and allocation trended higher (Fig. 3).

### Hypothesis 2b: Annual plants evolve drought escape strategies while perennials evolve drought tolerance under increasing drought stress

After subsetting the dataset to focus on plants exposed to drought (n=397), we found significant effects of trait category and the interaction between trait category and life history (QM=23.06, df = 2, p = 0.0003, Table S2). As expected, we detected evidence for the evolution of drought escape in annual descendants through earlier phenology (Fig. 4). Surprisingly, we also found evidence of earlier phenology in perennial plants as well, which is in opposition to our hypothesis that perennials will evolve drought avoidance/tolerance which would correspond with later flowering.

**Figure 4.**
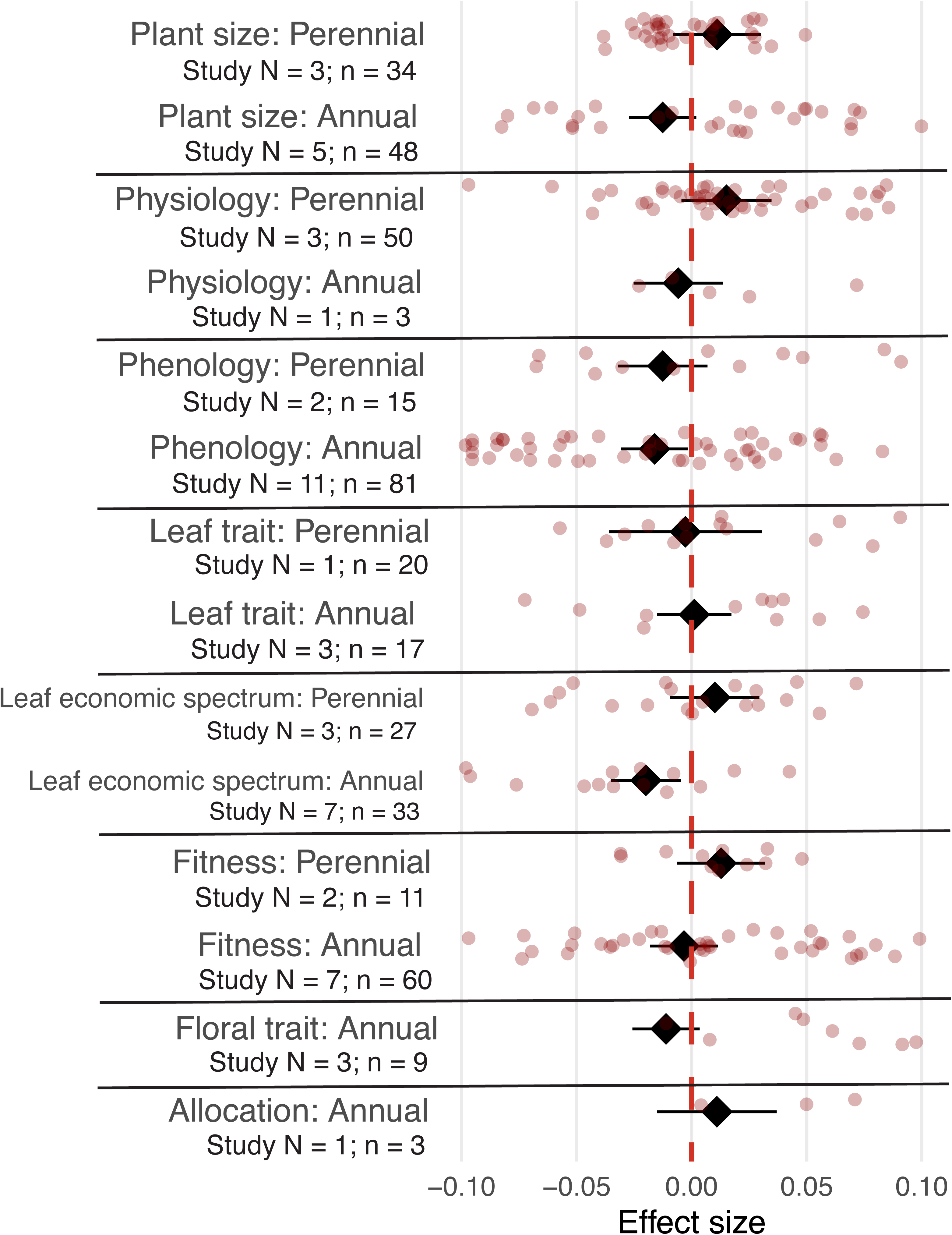
Interaction effects of trait category and life history on evolutionary rates for drought avoidance and escape trait data subset. Effect sizes are represented by black diamonds, with horizontal lines indicating 95% confidence intervals. Negative effect sizes would indicate shifts towards drought escape, while positive effect sizes would indicate shifts towards drought avoidance. Red points represent raw effect sizes. The x-axis is truncated to highlight patterns more clearly, full data is available as supplementary material. The overlap of confidence intervals with zero for some categories suggests non-significant effects in those cases. Study N indicates the number of studies that included each trait category; n is the number of observations included in the analysis. Dashed red vertical lines represent an effect size of 0.

### Hypothesis 3: Increasing environmental variability favors the evolution of phenotypic plasticity

After subsetting the dataset to just studies that imposed a drought treatment (n = 271), we calculated and analyzed the change in RDPI for ancestors and descendants and found significant effects of trait category (QM = 2235.7900, df = 7, p-val < .0001) and the interaction between trait category and life history (QM = 984.45, df = 4, p-val < .0001, Table S2). We found no trends in plasticity for perennial plants. For annual plants, we found an increase in plasticity for phenology and physiology traits, and a decrease in plasticity for LES traits (Fig. 5).

**Figure 5.**
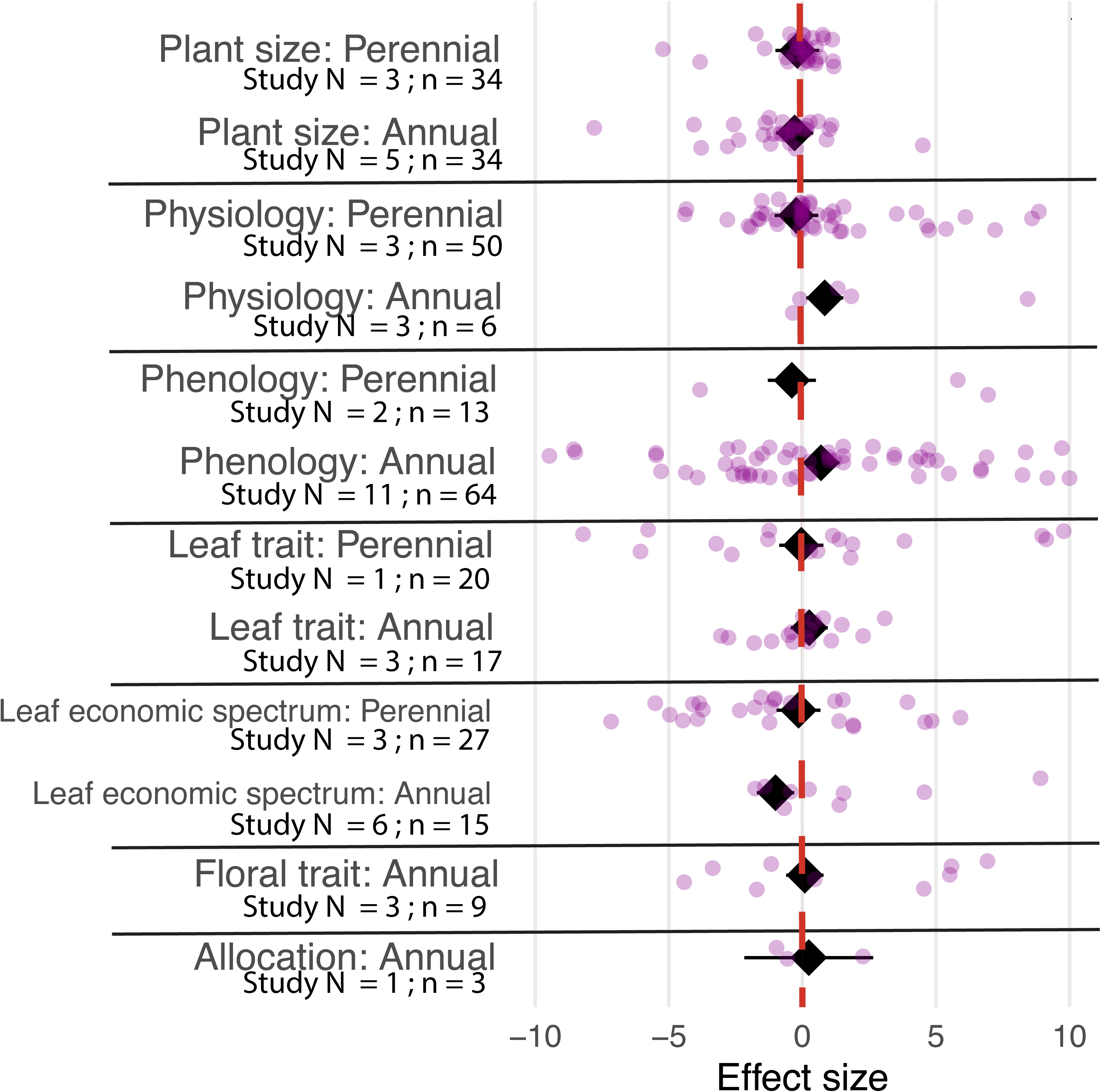
Interaction effects of trait category and life history on evolutionary rates of plasticity (as measured through the relative distance plasticity index) for drought avoidance and escape trait data subset. Effect sizes are represented by black diamonds, with horizontal lines indicating 95% confidence intervals. Negative effect sizes indicate reduced plasticity, while positive effect sizes would indicate an increase in plasticity. Purple points represent raw effect sizes. The x-axis is truncated to highlight patterns more clearly, full data is available as supplementary material. The overlap of confidence intervals with zero for some categories suggests non-significant effects in those cases. Study N indicates the number of studies that included each trait category; n is the number of observations included in the analysis. Dashed red vertical lines represent an effect size of 0.

## Discussion

Our review of resurrection studies revealed significant evolutionary changes in over 50% of observed traits, adding to a growing body of work showing that rapid evolution in natural populations is common (Sultan et al., 2013; Franks et al., 2016; Christie et al., 2023; White et al., 2023; Karitter et al., 2025). However, it is not ubiquitous, as nearly half of the traits we examined did not show evolution. Furthermore, we found limited evidence for consistent differences between ancestors and descendants across many trait categories, likely because trait evolution proceeded in different directions depending on species, populations and context. Below, we highlight opportunities for future research to clarify when and where rapid evolutionary is most likely.

### Hypothesis 1: Annual plants respond more rapidly to environmental change than perennials

Our findings partially support this hypothesis. As expected, annuals showed shifts toward earlier phenology and resource-acquisitive traits indicative of a stress escape strategy, and evidence for increased plasticity in phenology and physiology, potentially reflecting greater trait lability under heightened environmental variation. In contrast, evolutionary shifts were less evident in perennials, with earlier phenology only detected in response to drought. Due to shorter generation times, annuals may simply evolve faster in absolute time than perennials (Friedman, 2020), which is particularly relevant as climate change and extinctions unfold on absolute timescales. Our dataset included a good balance of life histories, but only one study (Everingham et al., 2021) examined woody species. This underrepresentation is notable given the ecological importance of long-lived plants in carbon storage, ecosystem stability, and biodiversity support (Pan et al., 2011; Betts et al., 2017), as well as their potential for adaptive genetic variation (Bisbing et al., 2021; Caignard et al., 2024). While such studies are challenging due to seed availability and timescales, resurrection studies in woody perennials are feasible with foresight and planning.

### Hypothesis 2 a & b: Annual plants differ from perennials in the direction of trait evolution and will evolve drought escape strategies, while perennials evolve drought tolerance

We hypothesized that global change would favor the evolution of trait values associated with stress escape in annuals and resource conservative trait values linked with stress tolerance in perennials. In line with classic theoretical frameworks (Levitt, 1972; Grime, 1977; Wright et al., 2004; Volaire, 2018), we further predicted that, under drought stress, annual species would evolve towards drought escape strategies (Fig. 1). Indeed, across analyses, we found that annuals evolved earlier phenology and more resource acquisitive leaf economic spectrum traits, though we detected no significant changes in physiology traits. In contrast, the full model revealed few consistent evolutionary trends in perennial species and, contrary to expectations, our drought-specific model did not show evidence for the evolution of drought tolerance or avoidance in perennials.

Our results are consistent with shifts toward earlier reproduction in many spring-flowering species observed globally (Parmesan and Yohe, 2003; Thackeray et al., 2016; Inouye, 2022). Phenology was the most consistently measured trait across resurrection studies (Fig. 2), with 76% of studies assessing reproductive timing, likely contributing to the strong signal detected in our meta-analysis. While we found evidence for drought escape in annuals, our results do not support a general shift towards drought tolerance in perennials. Temporal and spatial patterns may play a greater role than life history in shaping adaptive responses. For example, repeated late-season droughts could favor drought escape in both annuals and perennials (Heschel and Riginos, 2005).

One set of resurrection studies that illustrates this general trend of evolution toward stress escape and a resource acquisitive strategy was conducted with the annual plant *Brassica rapa*. These studies revealed adaptive evolution of earlier flowering following a multi-year late-season drought, consistent with escape (Franks et al., 2007; Weis et al., 2014). Rapid evolution, rather than plasticity, drove changes in traits like developmental timing and WUE (Franks and Weis, 2008; Franks, 2011b), with consistent patterns across populations, drought episodes, and in experimental evolution (Hamann et al., 2018; Johnson et al., 2022a). However, escape strategies showed limits under extreme drought (Hamann et al., 2018), underscoring the need to understand the boundaries of adaptation. Genomic analyses further revealed the genetic basis of these responses (Franks et al., 2016; Hamann et al., 2021c), highlighting the value of integrating genomic tools in future resurrection studies.

### Hypothesis 3: Increasing environmental variability favors the evolution of phenotypic plasticity

Increasingly variable conditions may impose novel selection on reaction norms (e.g., Nicotra et al., 2010; Chevin and Hoffmann, 2017). Our analysis detected increased plasticity in annual plants for phenology and physiology, and decreased plasticity for leaf economic traits. While plasticity can promote short-term persistence (Richardson et al., 2017) it may not suffice long term (Vinton et al., 2022) and is not always adaptive (Ghalambor et al., 2007; Duputié et al., 2015). Future resurrection studies should link reaction norms to fitness via selection analyses to assess adaptive value. Physiological traits like WUE and photosynthesis, which respond quickly and reversibly to environmental change (Grime and Mackey, 2002), may be especially prone to evolve plasticity under variability (Stotz et al., 2021). Yet few resurrection studies have quantified traits associated with physiology and resource allocation (Fig. 2). Nevertheless, drought can drive shifts in root traits and biomass allocation to enhance water acquisition (Padilla and Pugnaire, 2007; Comas et al., 2013; Lynch, 2013). CO_2_ enrichment, warming, and drought interact to exert selection on physiology, including WUE, stomatal conductance, and photosynthesis (Ainsworth and Rogers, 2007; Reich et al., 2018; Sendall et al., 2024). Future resurrection studies in variable environments could simultaneously assess evolutionary shifts in these traits and test for adaptive plasticity.

Reduced plasticity in leaf economic traits may reflect declining responsiveness to environmental variation (Stotz et al., 2022). Stress may promote stronger trait integration (Schlichting, 1986; Chapin et al., 1993; Pigliucci, 2005), constraining the evolution of plasticity. Geographic differences in climate change exposure (IPCC, 2023) may also matter: plasticity may not evolve if climatic variability remains unchanged. For example, Wooliver et al. (2020) found reduced thermal breadth in a southern *Mimulus cardinalis* population, likely due to decreased seasonality, and Branch et al. (2024) found plasticity loss in southern and gain in northern populations, aligning with spatial drought gradients. These findings emphasize the need to consider both mean climate shifts and variability over time. Experiments that expose ancestors and descendants from multiple populations to diverse environments can reveal how evolution, plasticity, and their interaction shape trait change (Anstett et al., 2021; Sheth et al., 2025), and can uncover the genetic basis of these responses (Franks et al., 2018).

### Rate of evolution in annual plants

Given that the generation time of annual species is known to be one year, we calculated the rate of evolution in haldanes from the studies included in our literature review. For these annual species, we found rapid rates of evolution, with the evolutionary rates of some trait categories reaching as high as 1.65 haldanes, which is substantially higher than in previous reviews (Hendry and Kinnison, 1999; Bone and Farres, 2001). These calculations add to a growing body of evidence that evolution in natural populations can be extremely rapid, particularly under strong selection.

We found extensive variation across trait categories in the rate of evolution, with leaf economical spectrum and phenological traits showing the fastest evolution, moderate rates of evolution for fitness and plant size, and slower rates for traits associated with resource allocation, floral characteristics and physiology. This result accords with prior studies that indicate that flowering time can evolve very rapidly and often, as we found, to earlier timing (Austen et al., 2017), while plant physiology traits are often highly plastic but may be somewhat less evolutionarily labile. It would be premature to draw conclusions from the seven reports of evolution of allocation-based traits, and the differences among trait categories should be interpreted with caution given the small sample sizes of some of these categories. However, these results highlight some potentially interesting trends and indicate that future resurrection experiments should attempt to quantify a greater breadth of phenotypes.

Rapid evolution can be consistent with adaptation, but it can also be caused by drift, especially in small populations, and it can be maladaptive, particularly under stress (Battlay et al., 2023; Fussmann and Kopp, 2023). On its own, the resurrection approach cannot distinguish adaptive from maladaptive or neutral evolution, but researchers can examine the fitness consequences of trait shifts (Hamann et al., 2018). Future studies that expose lineages to ancestral and contemporary environments can test whether selection favors ancestral trait values in environments that reflect historical climates and descendant trait values in environments that reflect contemporary climates. Furthermore, exciting quantitative genetic experiments could be designed to contrast the adaptive potential of ancestral and descendant lines through estimates of genetic variance in fitness and traits under different climate scenarios.

### Constraints on evolutionary rescue

Notably, nearly half of the examined cases did not document significant trait differences between ancestral and descendant generations, highlighting the challenges of accurately predicting the likelihood of evolutionary rescue. A key area for future research is further disentangling when, why, and where rapid evolutionary responses are most likely. Theory and empirical results from microcosm experiments suggest that evolutionary rescue is most likely when population sizes are large, standing genetic variation is high, and the rate of environmental change is gradual (Carlson et al., 2014). Integrating resurrection studies with data on demography (e.g., Sheth and Angert, 2018), quantitative genetic variation, climate anomalies, and knowledge of the agents and targets of selection (Wadgymar et al., 2022) will inform predictions of when and how evolution could enable or hinder long-term population persistence with climate change. As we continue to broaden the taxonomic and geographic breadth of resurrection research and expand the spatial and temporal sampling within these studies, our understanding of the contexts in which populations are most likely to adapt to climate change will improve.

### Recommendations for future studies

Few resurrection studies have examined biotic interactions, invasive species, mating system evolution, multiple environmental factors, or range-wide variation. Here we describe progress toward evaluating how complex suites of changing abiotic and biotic environments influence plant evolution.

#### Biotic interactions

We found only three studies that examined evolution in response to herbivory (Beaton et al., 2011; Bustos-Segura et al., 2014; Rauschkolb et al., 2022b) and one that measured fungal pathogen susceptibility between ancestral and descendant generations (O’Hara et al., 2016). Novel communities, including changing pollinators, seed dispersers, herbivores, and competitors, could influence the evolutionary trajectory of plants (Williams and Jackson, 2007; Alexander et al., 2015; Lancaster et al., 2017; Miller et al., 2023). Furthermore, climate change can lead to phenological mismatches between interacting species—such as plants flowering earlier than their pollinators (Rafferty et al., 2015; Visser and Gienapp, 2019). In addition, anthropogenic stressors like habitat loss (Spiesman and Inouye, 2013) and pesticides or herbicides (Ramos et al., 2023) can further disrupt biotic interactions. Climate change directly influences the abundance and activity patterns of the species that interact with plants, such as pollinators, seed dispersers, herbivores, soil microbes, and competitors (Mokany et al., 2014; Rafferty, 2017; Hamann et al., 2021a). In addition, climate change shifts key plant traits that have evolved in response to selection imposed by these mutualists and antagonists (Hamann et al., 2021b). Other changing biotic interactions—such as increased herbivory (Hamann et al., 2021a), altered microbial symbioses (Compant et al., 2010), and disrupted seed dispersal (Mokany et al., 2014)—represent promising areas for resurrection studies.

The resurrection approach can also shed light on long-standing questions in invasion biology. For example, resurrection studies could be used to test the hypothesis that invasive plants may show stronger evolutionary response than native plants, as faster adaptation often strengthens invasibility (Novak, 2007; Prentis et al., 2009). Indeed, Beaton et al. (2011) tested the evolution of increased competitive ability (EICA; Blossey and Notzold, 1995) hypothesis by comparing ancestral seeds of *Lespedeza cuneata* (Fabaceae) collected in the native range to descendants grown in both the native and invasive ranges. They found that invasive descendants exhibited greater competitive ability and reduced investment in herbivore defenses, representing a powerful test of the EICA hypothesis. Given the strong effects of invasive species on native biodiversity (Mollot et al., 2017), it is crucial to examine whether novel selection will increase (Blair and Wolfe, 2004; Woods and Sultan, 2022) or decrease (Colautti et al., 2010; Felker-Quinn et al., 2013) the fitness of these species.

#### Mating system evolution

Climate change could alter the evolution of mating systems (Eckert et al., 2010; Jones et al., 2013; Ramos and Schiestl, 2019; Hamann et al., 2021c; Rusman et al., 2025; Traine et al., 2024), yet our meta-analysis did not detect signatures of shifts in floral traits consistent with mating system evolution. Increasing drought stress and global reductions in pollinator abundance (Eckert et al., 2010; Jones et al., 2013; Cornelisse et al., 2025) could favor the evolution of traits that enable selfing, such as reduced herkogamy (spatial separation between anthers and stigma) and dichogamy (temporal separation between anthers and stigma), and the associated floral traits, like reduced floral rewards and flower sizes. For example, pollinator decline in a native population of *Viola arvensis* (Violaceae) was associated with smaller flowers and increased selfing in the descendant generation (Cheptou et al., 2022). In contrast, Thomann et al. (2015a, 2015b) observed increased flower size in native populations of *Adonis annua* (Asteraceae) and *Centaurea cyanus* (Ranunculaceae) under pollinator decline, which could arise to enhance pollinator attraction. Thus changes in pollinator activity can drive evolutionary responses in native plant populations (Traine et al., 2024; Rusman et al., 2025). Resurrection studies provide a unique opportunity to test whether selfing could evolve as a mechanism for reproductive assurance. Future resurrection studies using congenic pairs of species that differ in mating system could directly evaluate the role of this key reproductive system in rapid evolutionary dynamics under climate change.

#### Multifactorial approaches

Many published resurrection experiments (37.5% of all studies) to-date have focused on drought, either through experimental manipulation or by collecting seeds before and after severe droughts. Indeed, drought was the most common manipulation in the reviewed resurrection studies, followed by heat stress, while fewer studies altered light, herbivory, pathogens, or competition (Table S1). Climate change is inherently multifactorial, simultaneously altering temperature, precipitation, nutrient cycles, season length, and biotic interactions, and increasing the frequency and severity of extreme events, and even increased precipitation in many regions (IPCC 2023). Although most experiments tested single factors, 19% included multiple global change drivers, and we encourage future explorations of multiple aspects of climate change (e.g., Wooliver et al., 2020; Preston et al., 2022; Albano et al., 2025). Finally, we found that 72% of resurrection studies were conducted in controlled environments. When specific selective agents are unknown or too complex to simulate in the lab, field gardens offer a way to measure adaptation under realistic conditions (Karitter et al., 2025; Sheth et al., 2025).

#### Examination of multiple traits

While prior work has shown that populations can evolve rapidly in response to environmental change (Anstett et al., 2021; Franks et al., 2007; Hamann et al., 2018), little is known about how such adaptations affect trait syndromes, trade-offs, and potential fitness costs. Adaptation to one stressor may compromise performance under others (Hereford, 2009), shifting ecological strategies in ways that carry long-term consequences (Hamann et al., 2021a). For instance, does increased drought tolerance reduce competitive ability? Do stress-adapted perennials evolve faster life histories due to increasing costs of reproduction under stress (Hamann et al., 2021d)? Additionally, exposing ancestral and descendant lines to novel environments may reveal cryptic genetic variation—unexpressed under past conditions—that signals hidden adaptive potential (Paaby and Rockman, 2014; Josephs, 2018). We advocate for multivariate and multifactorial resurrection studies to capture the full range of trait covariation and trade-offs across diverse environmental conditions to test how ecological strategies and trait syndromes are evolving, and to examine constraints on adaptation.

#### Range-wide studies

Since life history, demography, climate anomalies, phenotypic selection, and genetic variation often vary across spatial and environmental gradients (Kinnison and Hairston, 2007; Sexton et al., 2009; Kokko et al., 2017; Pironon et al., 2017), populations across a species’ range could differ substantially in their rates of evolutionary responses to selection (Sheth and Angert, 2018). Even the earliest resurrection study involving flowering plants considered that evolutionary responses could depend on the environmental context and thus background selection that a population has experienced (Franks et al., 2007, reviewed in Franks et al., 2018). Franks et al. (2007) included populations from a wet and dry site and documented greater evolutionary responses to a period of drought in the wet-origin population. More recent resurrection studies have expanded to include populations that span species’ ranges, particularly studies of *Mimulus* (Phrymaceae) species: *Mimulus guttatus* (37 populations; (Kooyers et al., 2021), *Mimulus laciniatus* (9 populations; Dickman et al., 2019), and *Mimulus cardinalis* (6-11 populations per study; Anstett et al., 2021; Vtipil and Sheth, 2020; Branch et al., 2024).

Such studies allow for robust tests of geographic variation in response to change. For instance, Kooyers et al. (2020) measured trait variation in multiple populations of *M. guttatus* that were experiencing varying drought intensity and found inter- and intra-population variation in phenology, flowering node, and height at first flower. Populations did not show predictable patterns of evolution, but the highest magnitude changes were associated with populations that had greater heritable variation in the ancestral generation, highlighting the importance of standing genetic variation than drought intensity. Other studies have included large numbers of populations spanning broad environmental gradients. For example, Al-Hajaj et al (2022) sampled *Hordeum spontaneum* (Poaceae) from across the entirety of Jordan, with populations from five distinct bioclimatic zones, and detected population- and region-level differences between and amongst generations, with flowering time shifting earlier in response to changes in climate in the regions experiencing the warmest climates and strongest shifts in climate.

Populations can vary in response to change due to a multitude of factors including local climate, population size, history, and isolation. Further, more work is needed to understand range-wide trends of adaptability and whether range edges are in danger of extirpation due to an inability to adapt (Hampe and Petit, 2005; Pennington et al., 2021). For example, researchers could bank seeds from natural populations in the trailing (warming) edge, the center, and the leading (historically cooler) edge of the geographic distribution. Studies that expose ancestors and descendant lineages from populations distributed across the range to multiple environmental conditions could evaluate the rate of evolution, whether that evolution was adaptive, and to what extent the adaptive potential varies across the landscape. Furthermore, these studies could examine how different demographic parameters influence the evolutionary trajectory of populations, and whether strong selection imposed by climate change decreases standing genetic variation in populations, thereby reducing the capacity for future adaptive evolution.

### Additional opportunities and resources for future studies

Our literature review revealed numerous opportunities for continued application of resurrection studies. One key resource available for studies in North America is *Project Baseline* (Franks et al., 2008; Etterson et al., 2016), which established long-term seed storage of a set of focal species, supporting research proposals that utilize these collections. Further, conservation seed bank collections have been used in resurrection studies (e.g. Everingham et al., 2021) and can be considered for future resurrection attempts (Mattana et al., 2025). In addition, we recommend that future resurrection studies include locations outside of North America and Europe, which are overrepresented in the database of resurrection studies. Finally, genomic studies thus far have largely confirmed the genetic basis of phenotypic changes. Challenges remain, such as the often-limited ability to link genomic changes to phenotypic outcomes (Hoban et al., 2016) but combining multi-generational and resurrection studies with whole-genome sequencing has the potential to greatly enhance our understanding of plant adaptation (Brown and Koenig, 2022; Anstett et al., 2024).

## Conclusions

Resurrection studies have deepened our understanding of the evolutionary capacity of species under global change. From an analysis of 40 studies conducted during the past 20 years, we detected widespread rapid evolution in natural plant populations, including the evolution of drought escape in annual plants, which matches evolutionary theory. However, it is also notable that nearly half of the examined cases did not document significant trait differences between ancestral and descendant generations, highlighting the challenges of accurately predicting the probability of evolutionary rescue. One critical area for future research is assessing whether rapid, directional selection across generations results in a reduction of genetic variance or an increase in genetic correlations opposing selection, potentially constraining future adaptive potential (Kopp and Matuszewski, 2014). Additionally, most resurrection studies have focused on single stressors, yet natural environments are inherently multifactorial. Investigating potential fitness trade-offs—such as whether adaptation to drought escape compromises competitive ability or reduces resistance to herbivory—will provide a more realistic picture of how plants evolve under simultaneous pressures. Combining resurrection studies with reciprocal transplants in the field, where multiple fitness components and their tradeoffs can be quantified in the presence of relevant biotic interactions, is a particularly powerful approach. By moving toward more integrative, multivariate approaches and including a broader range of life histories, resurrection studies can play a key role in predicting and supporting species persistence under accelerating global change.

## Acknowledgements

We thank the many researchers who conducted the resurrection experiments that informed this study. We would also like to thank Craig Osenberg for guidance on details of the meta-analysis. This work was supported by grants from the National Science Foundation (IOS-2220927 and DEB-1553408 to J. Anderson, DEB-2131815 and IOS-2220928 to S. Sheth, PRFB-2305903 to L. Pennington, and IOS-1546218 to S. Franks). S. Sheth was also supported by a Research Capacity Fund (HATCH) project award no. 7002993 from the U.S. Department of Agriculture’s National Institute of Food and Agriculture.

---

Table S1. Summary table of resurrection studies. NR = Not reported. An asterisk in the life history column indicates that life history was not reported in text and we searched for accurate life history information for that species. Mating system reported as in text. An asterisk in ‘Was the evolutionary change adaptive?’ column indicates the change was discussed as adaptive but was not formally tested as having resulted in increased fitness.
[Table attached as excel sheet]

**Table S2:** Results from the meta-analysis mixed model for the full model (H1, H2a), drought analysis (H2b), and plasticity analysis (H3). QM model statistic, df, and p value reported.

|  | Fixed effect | QM | df | p |
| --- | --- | --- | --- | --- |
| Full Model | Life history | 1.1 | 2 | 0.58 |
| n = 1746 | Trait category | 133.12 | 8 | < **0.0001** |
|  | Life history * Trait category | 111.94 | 6 | < **0.0001** |
| Drought analysis | Life history | 0.55 | 2 | 0.76 |
| n = 397 | Trait category | 98.19 | 8 | < **0.0001** |
|  | Life history * Trait category | 23.06 | 5 | **0.0003** |
| Plasticity analysis | Life history | 0.11 | 2 | 0.95 |
| n = 271 | Trait category | 2235.8 | 7 | < **0.0001** |
|  | Life history * Trait category | 984.45 | 4 | < **0.0001** |

**Table S3:**
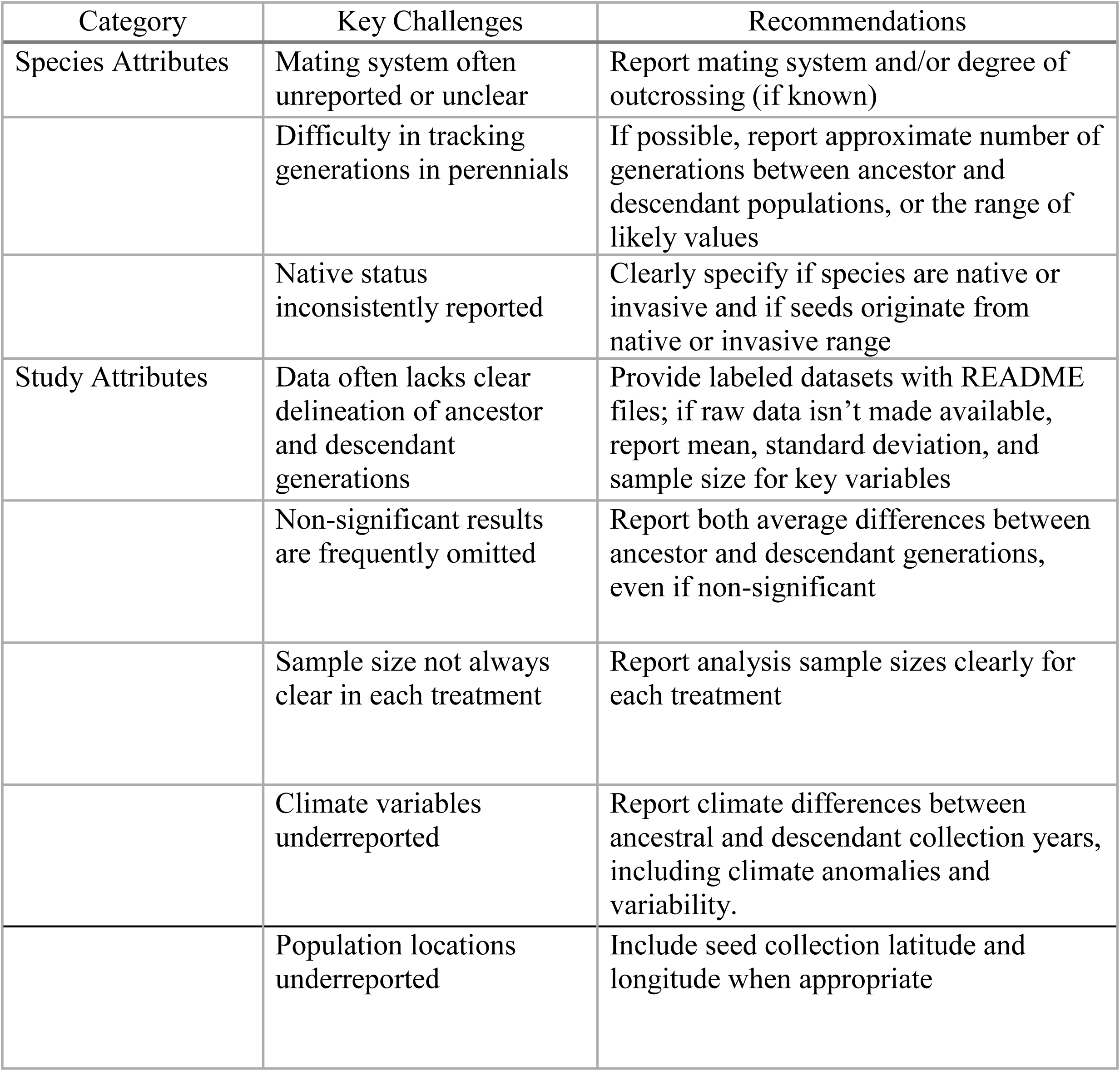
We encountered several challenges during qualitative and quantitative data extraction that could likely be alleviated by more standardized reporting of key details in resurrection studies, though such challenges are common in meta-analyses (Koricheva & Gurevitch 2014). Here, we summarize key challenges we encountered and recommendations for reporting in future resurrection studies.

| Category | Key Challenges | Recommendations |
| --- | --- | --- |
| Species Attributes | Mating system often unreported or unclear | Report mating system and/or degree of outcrossing (if known) |
|  | Difficulty in tracking generations in perennials | If possible, report approximate number of generations between ancestor and descendant populations, or the range of likely values |
|  | Native status inconsistently reported | Clearly specify if species are native or invasive and if seeds originate from native or invasive range |
| Study Attributes | Data often lacks clear delineation of ancestor and descendant generations | Provide labeled datasets with README files; if raw data isn't made available, report mean, standard deviation, and sample size for key variables |
|  | Non-significant results are frequently omitted | Report both average differences between ancestor and descendant generations, even if non-significant |
|  | Sample size not always clear in each treatment | Report analysis sample sizes clearly for each treatment |
|  | Climate variables underreported | Report climate differences between ancestral and descendant collection years, including climate anomalies and variability. |
|  | Population locations underreported | Include seed collection latitude and longitude when appropriate |

**Figure S1:**
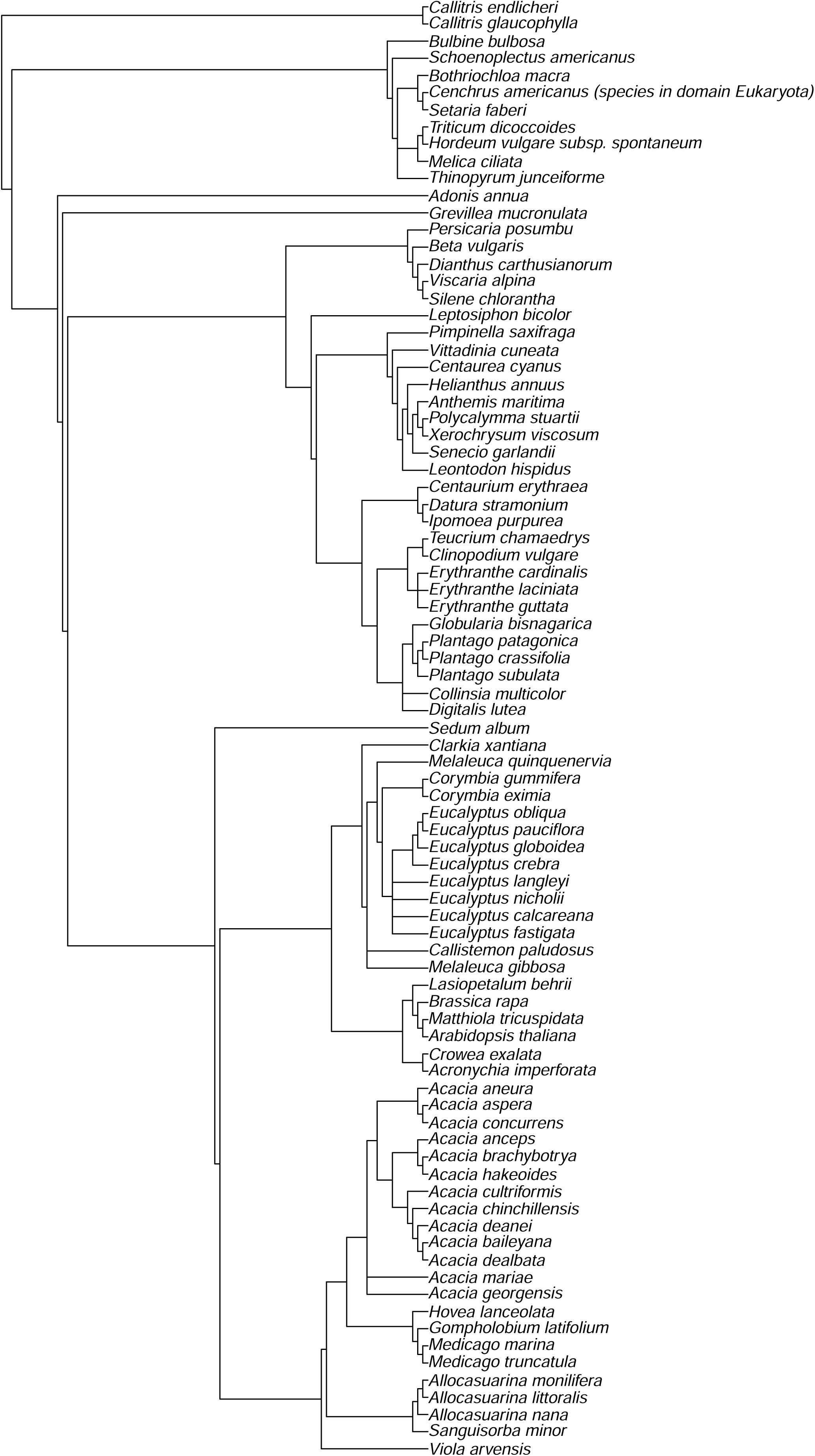
Phylogeny of species included in the formal meta-analysis

**Figure S2:**
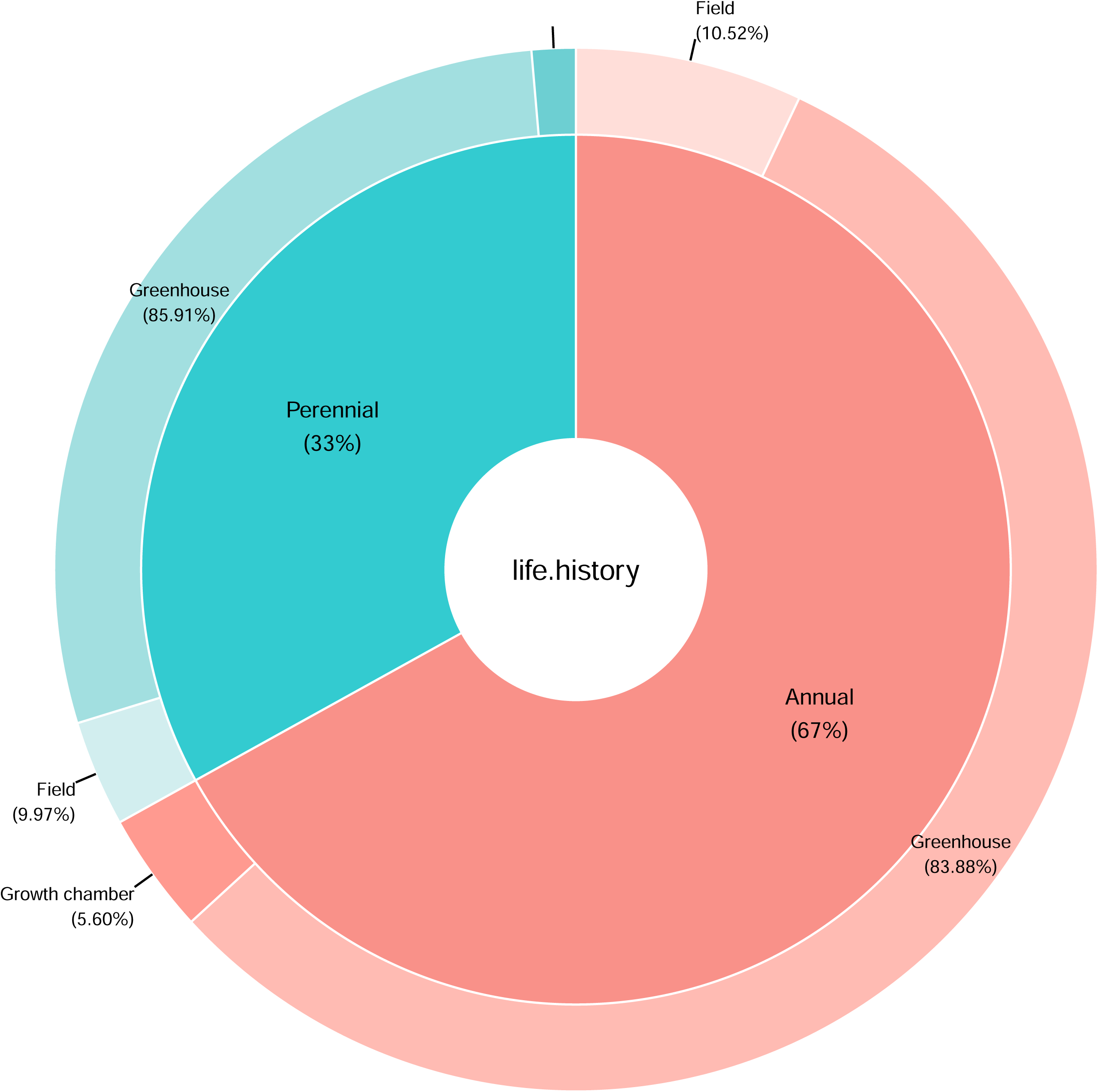
Percentage of observations for each experimental context for each life history category

**Figure S3:**
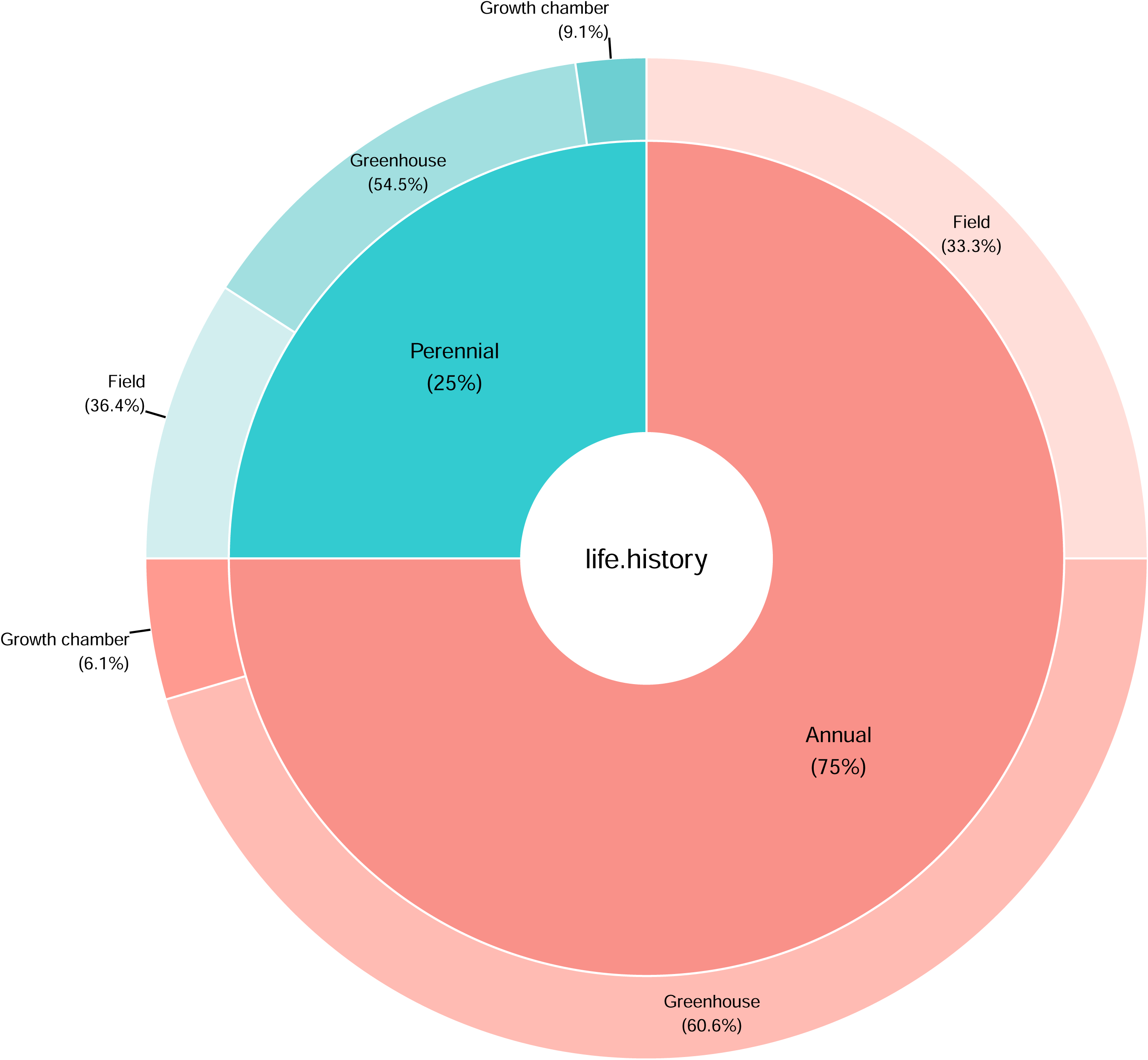
Percentage of studies for each experimental context for each life history category

**Figure S4:**
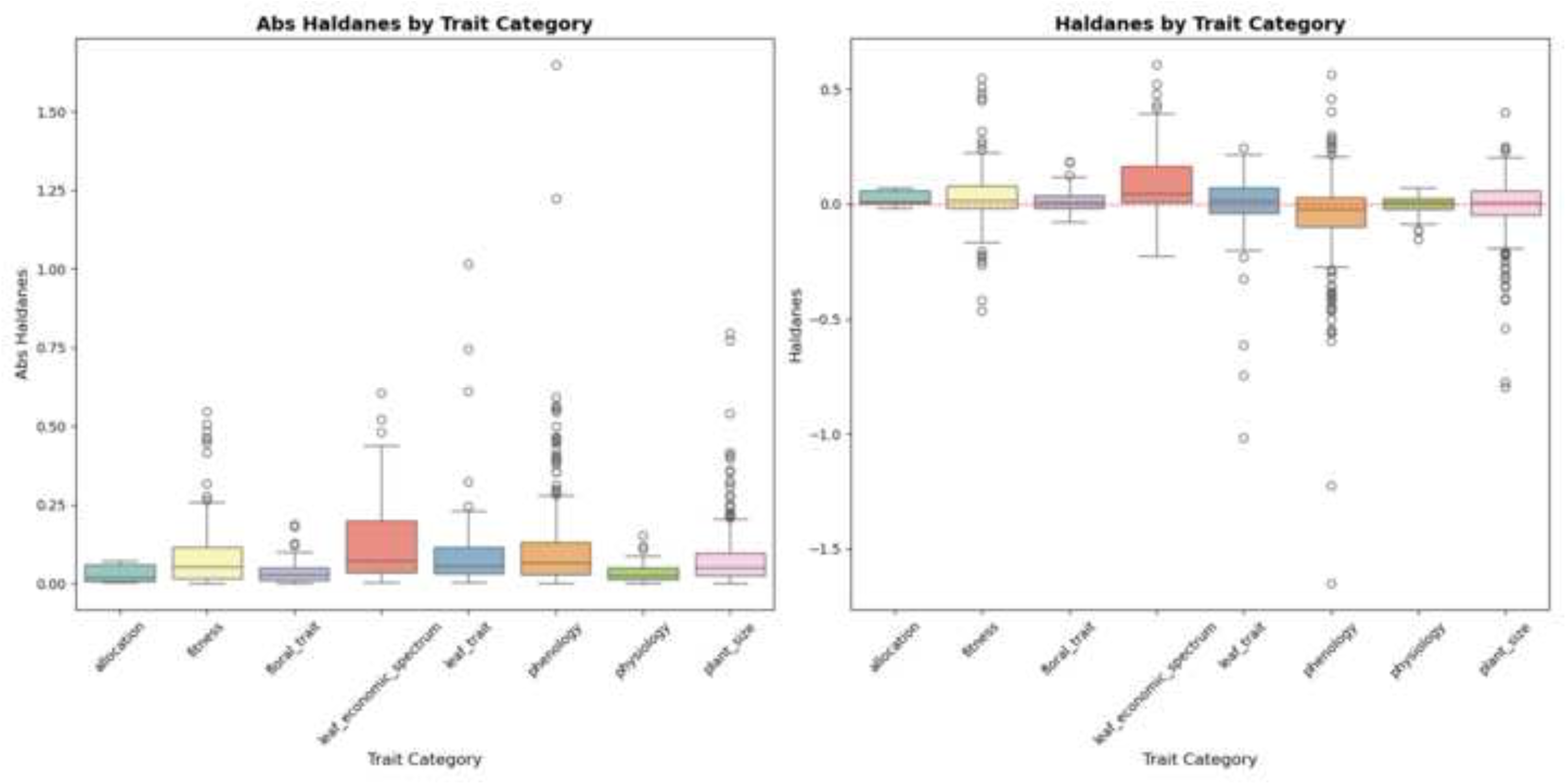
Boxplots of the absolute value of haldanes (left) and haldanes (right), reported by trait category.

**Figure S5:.**
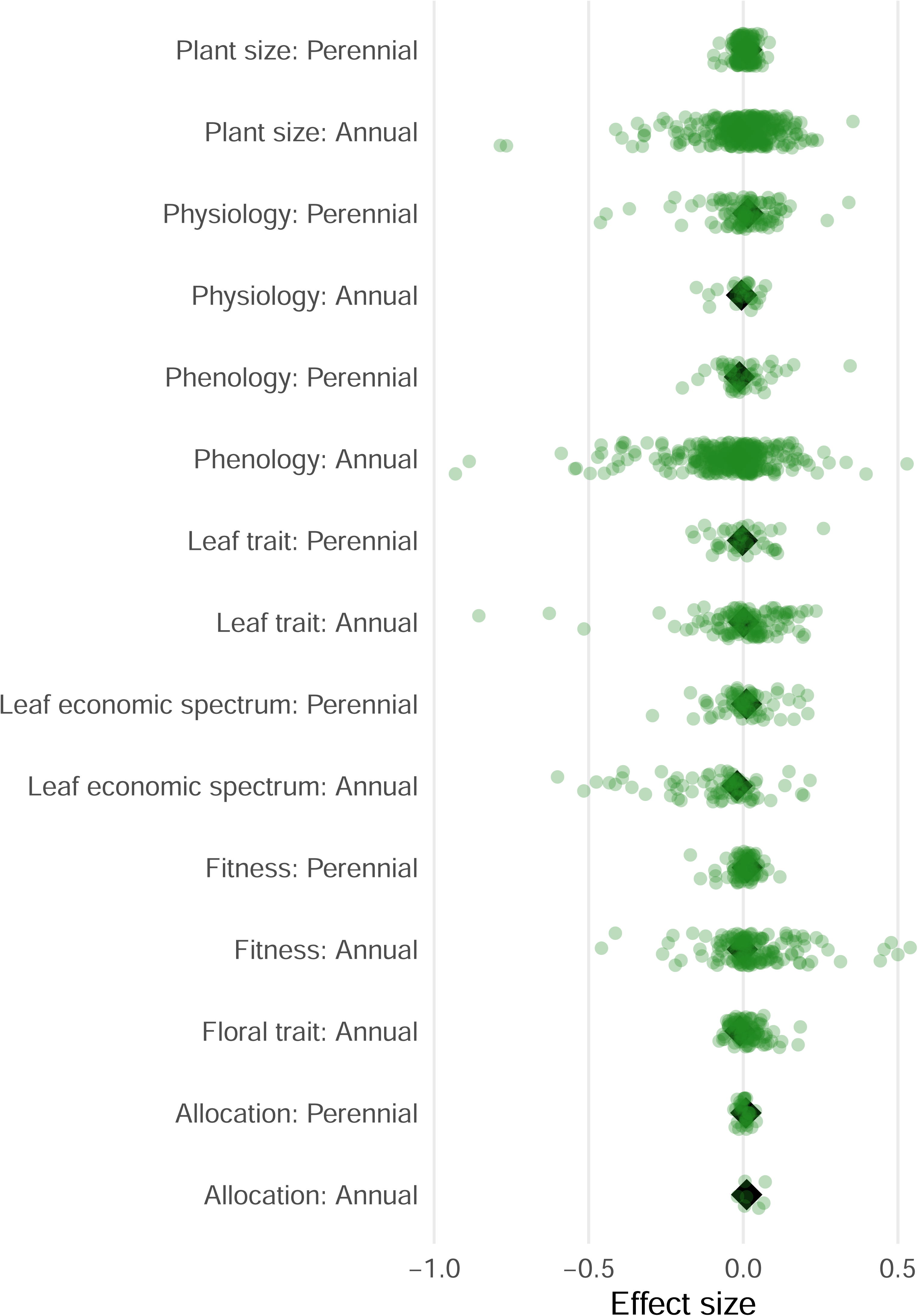
Interaction effects of trait category and life history on evolutionary rates. Effect sizes are represented by black diamonds, with horizontal lines indicating 95% confidence intervals. Positive values indicate that descendants have larger trait values than ancestors. Green points represent raw effect sizes; all data shown here for figure 3.

**Figure S6:.**
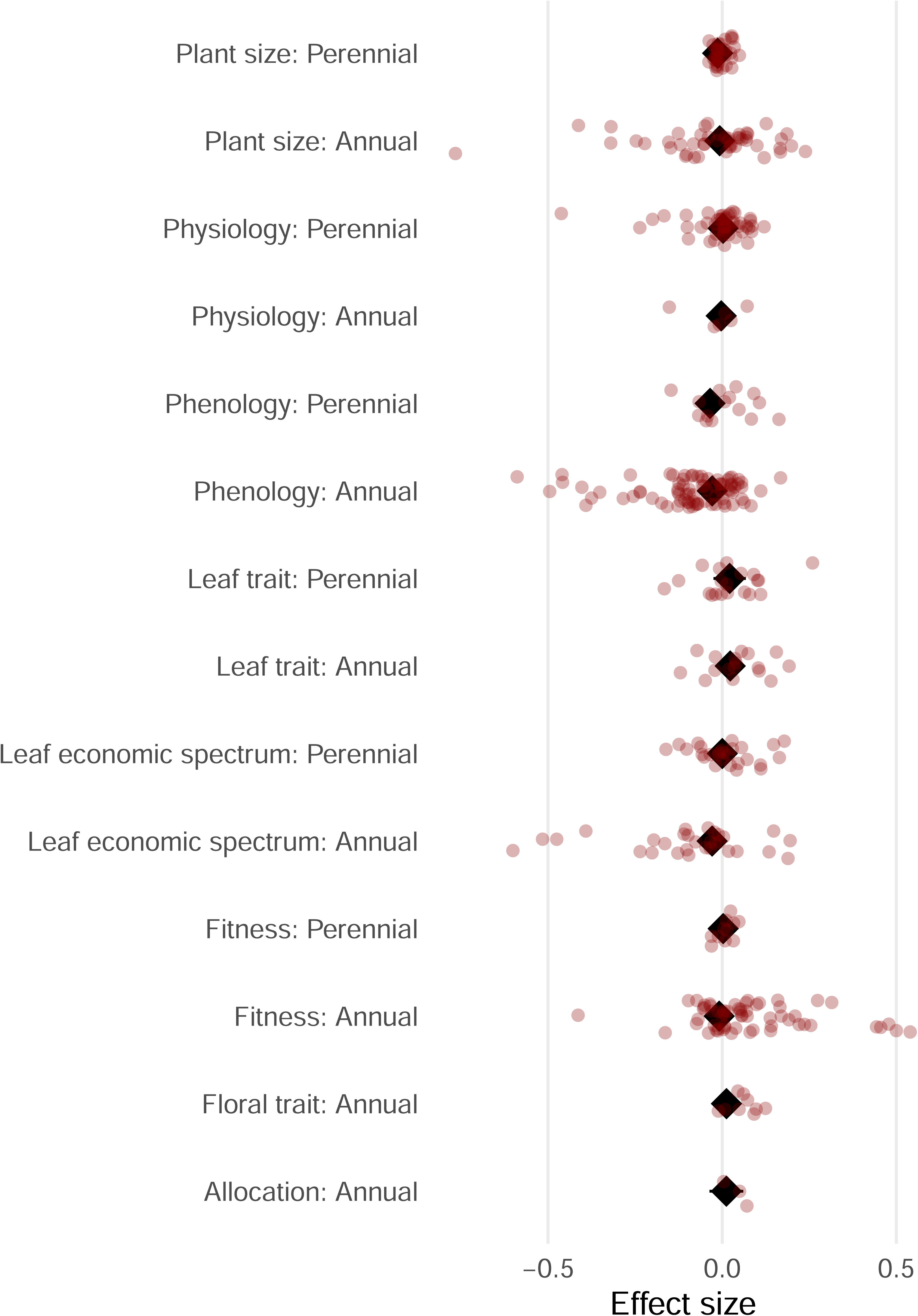
Interaction effects of trait category and life history on evolutionary rates. Effect sizes are represented by black diamonds, with horizontal lines indicating 95% confidence intervals. Positive values indicate that descendants have larger trait values than ancestors. Red points represent raw effect sizes; all data shown here for figure 4.

**Figure S7:.**
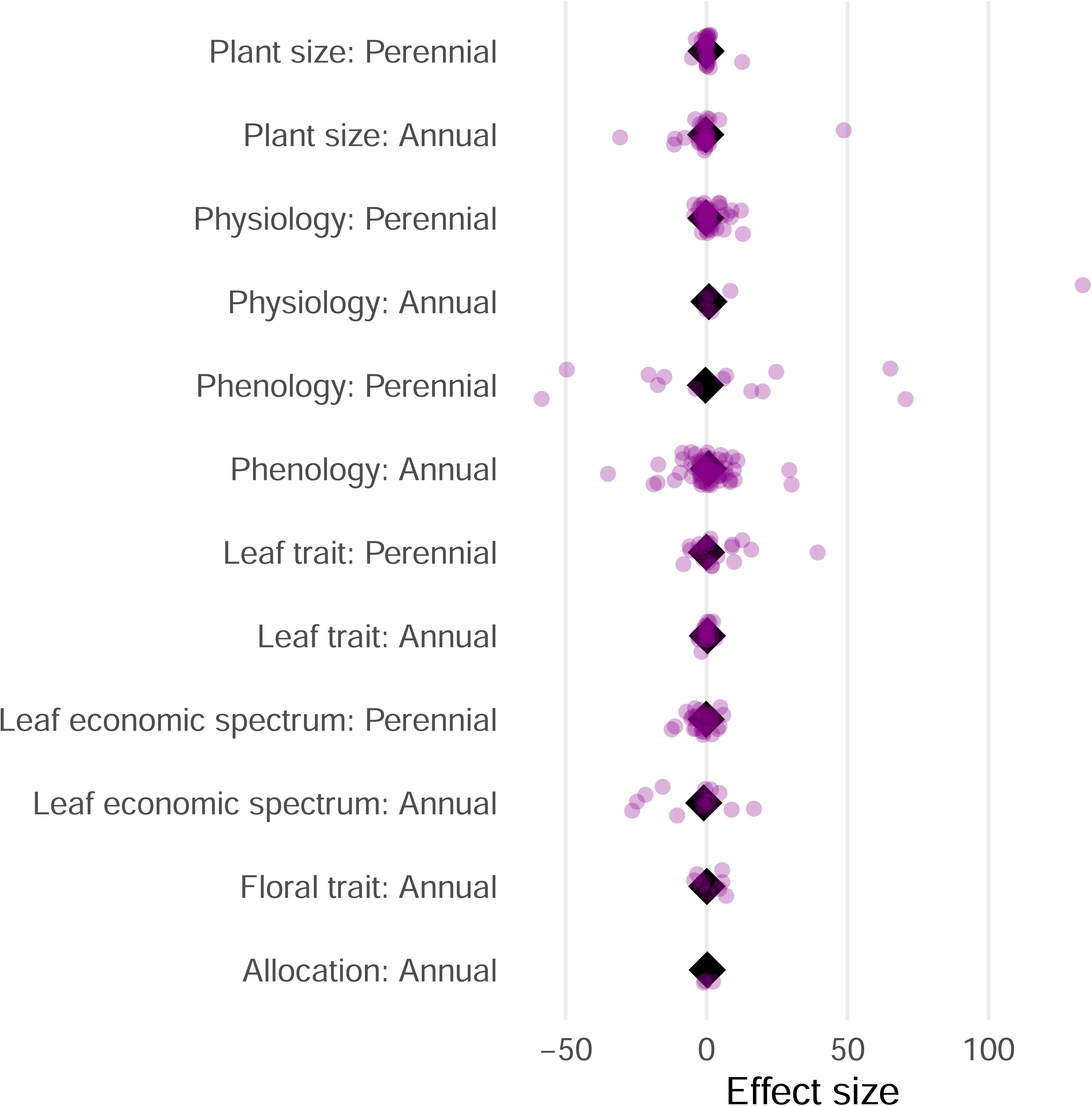
Interaction effects of trait category and life history on evolutionary rates. Effect sizes are represented by black diamonds, with horizontal lines indicating 95% confidence intervals. Positive values indicate that descendants have larger trait values than ancestors. Purple points represent raw effect sizes; all data shown here for figure 5.

